# Genetic and pharmacologic modulation of RAGE rescues the diabetes-mediated impairments to bone at multiple length scales

**DOI:** 10.64898/2026.04.02.716153

**Authors:** Kaitlyn S. Broz, Timothy Hung, Remy E. Walk, Suzanne LoTempio, Katharine M. Flores, Simon Y. Tang

**Affiliations:** Institute of Materials Science and Engineering, Washington University in St Louis, MO; Department of Orthopedic Surgery, Washington University School of Medicine, St Louis, MO; Department of Biomedical Engineering, Washington University in St Louis, MO

## Abstract

The bone matrix is precisely maintained and optimized to resist fractures. However, aging and disease deteriorate the bone matrix and increase fragility. Individuals with type 2 diabetes (T2D) have an elevated risk of bone fracture despite apparently normal bone mass. The chronic hyperglycemia in T2D promotes the formation of advanced glycation end-products (AGEs) in the bone tissue and modify the matrix mechanics. AGEs also bind to its receptor, RAGE, to activate inflammation and alter homeostasis. Using a leptin-receptor deficient mouse model of diabetes, we used a combination of high-resolution methods across multiple scales to evaluate the microarchitectural-, material- and cellular- level changes affected by the modulation of RAGE. To demonstrate the relevance of RAGE, we genetically ablated RAGE (RAGE-null) before the onset of diabetes; and to demonstrate the potency of RAGE as a disease modifying therapy, a RAGE antagonist (FPS-ZM1) was administered after prolonged diabetes. Diabetes impaired bone microstructure, the homeostatic actions of bone cells, the bone matrix nanomechanics, and whole-bone strength. The constitutive ablation of RAGE in diabetic animals prevented AGEs accumulation and the decline of trabecular connectivity; protected against the loss of osteocyte lacunae density and morphology; and maintained the matrix nanomechanics and bone strength. The inhibition of RAGE after the onset of diabetes reversed AGE accumulation and loss of bone volume; rescued osteocyte lacunae density and osteoclast activity; and restored matrix nanomechanics and bone strength. These results suggest that RAGE is a viable therapeutic target for diabetes-mediated impairments of bone quality.

**Graphical Abstract:** 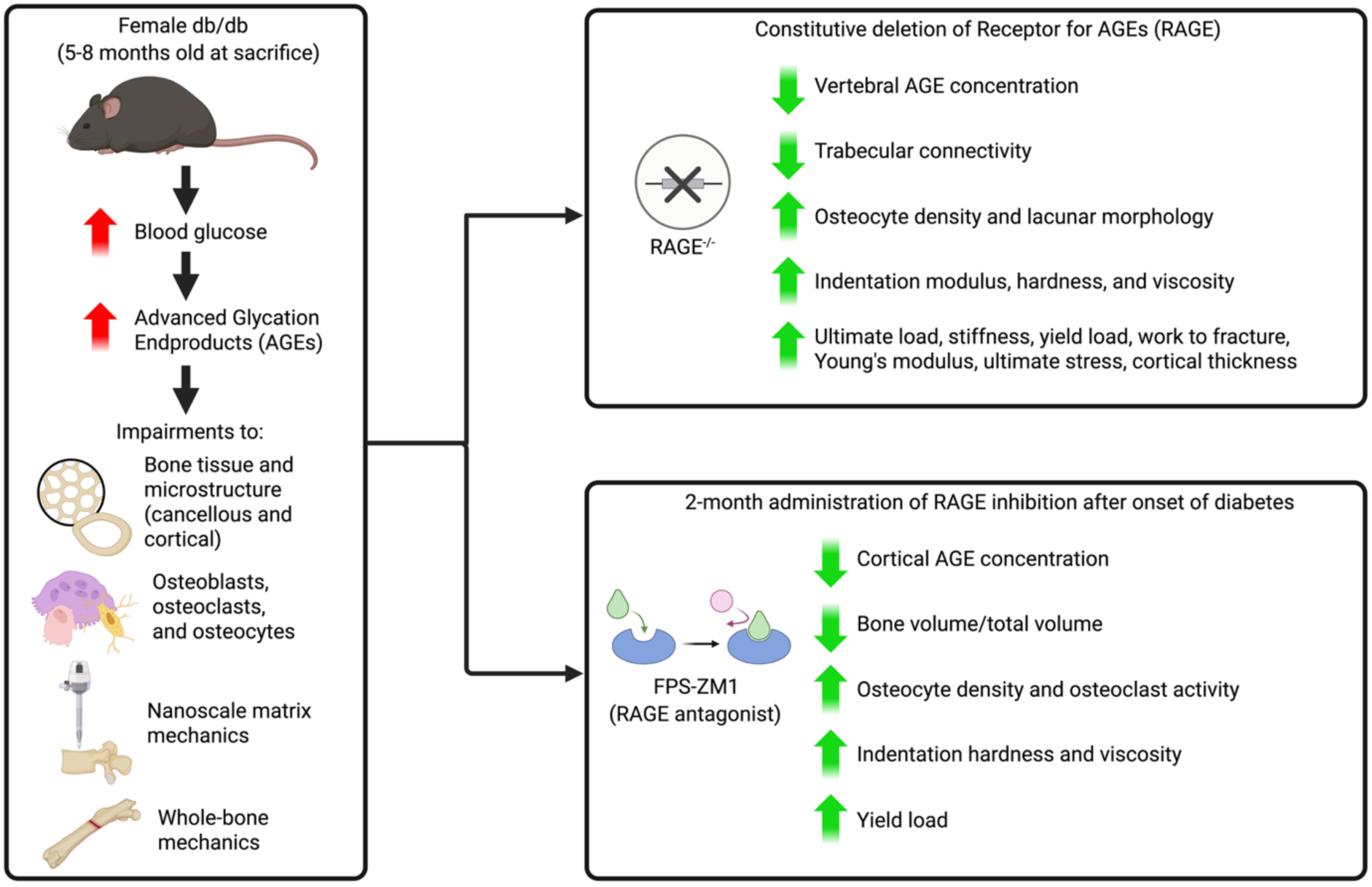

## 1. Introduction

Patients with type 2 diabetes (T2D) live with comorbidities that greatly impair their quality of life. The global health burden of T2D is rising due to the increasing incidence of T2D in the global population, which is expected to reach 9.5% by the year 2025.^1^ Comorbidities include vascular complications, neuropathy, and a number of musculoskeletal complications including increased fracture incidence.^2^ The incidence of vertebral fractures occur with 3.1x greater incidence in post-menopausal women with T2D compared to age matched controls.^3^ Yet, identifying the patients with T2D that are more likely to experience a bone fracture is quite difficult. Bone mineral density (BMD) does not appear to predict fracture risk in patients with T2D.^4^ Counterintuitively, patients with T2D present with a higher BMD compared to non-diabetic controls,^4^ and within the T2D population BMD is discordant with relative fracture risk.^4^ These studies reveal that there are impairments in the bone matrix of patients with T2D leading to the mechanical impairments and increased fracture risk.

The accumulation of advanced glycation endproducts (AGEs) has been widely recognized as a contributor to musculoskeletal complications in patients with T2D.^5,6^ AGEs are formed via the non-enzymatic glycation of amino acids and reducing sugars, also known as the Maillard reaction; AGEs tend to accumulate in high glucose environments like T2D.^5^ The abundance of bone’s organic matrix thus make it particularly susceptible to the formation of AGEs in T2D. The formation of AGEs in the bone’s organic matrix often crosslinks the protein components, and the accumulation of AGEs directly modify the matrix mechanics and fracture resistance of skeletal tissues.^6–8^ AGEs can also bind to their receptor, the receptor for advanced glycation endproducts (RAGE), and the activation of RAGE in bone cells affect their survival and function.^9–20^ Osteoclast activity, osteoblast activity, and osteocyte density are responsive to RAGE signaling. RAGE activation decreased osteoclast differentiation *in vitro*^9^ but divergent effects *in vivo*^10–12^. Osteoblasts and osteocytes exhibit decreased differentiation and increased apoptosis in high AGE and high glucose environments *in vitro*.^13–15^ Consistent with these observations, *in vivo* studies have also shown decreased osteoblast activity^16,17^ and reduced osteocyte density or impairments to osteocyte lacunar network^18–20^.

RAGE signaling is chronically elevated in patients with T2D^21,22^, and is upstream of the NF- κB inflammatory pathway^23^. Therefore the sustained activation of RAGE ineffectuates bone cell function in highly inflammatory environments like T2D.^24–26^ These disruptions to bone cell activity are associated with declines in whole-bone strength. Prior studies have shown that RAGE- deficiency (RAGE^-/-^) improves BMD and whole bone strength in db/db mice.^27,28^ However, how RAGE signaling affect in vivo bone remodeling dynamics, matrix nanomechanics, and whole bone strength remains unclear. Moreover, no studies have evaluated the ability of RAGE inhibition to rescue bone impairments after prolonged diabetes.

To demonstrate the clinical utility of targeting RAGE, we assessed the bone matrix mechanics, bone cell activity, and strength in diabetic mice following either preventative genetic ablation or therapeutic pharmacological inhibition. Our multi-scale evaluation reveals that ablating RAGE prevents diabetes-mediated declines in trabecular connectivity and osteocyte integrity, while the inhibition of RAGE after the onset of diabetes successfully restores bone volume and strength. Thus, we propose RAGE as a viable therapeutic target for reversing diabetes-mediated skeletal impairment.

## 2. Materials and Methods

All studies were completed with WUSM IACUC approval. The leptin receptor deficient (Lepr^db/db^; aka db/db) mouse were used in these studies. The lack of leptin-signaling due to the defective leptin receptor results hyperphagia that culminates in elevated morbid obesity, hyperglycemia, and insulinemia as early as 3-4 weeks of age.^29,30^ These db/db animals exhibit impaired fracture resistance as early as 3 months of age.^28^

### 2.1 Genetic ablation of RAGE

To perform genetic ablation of RAGE, we crossed C57BL/6 mice carrying a defective RAGE allele with the db/db animals.^31,32^ From these crosses, we allocated 6-7 animals into each group (n=6-7) wt (RAGE^wt/wt^; Lepr^wt/wt^), db/db (RAGE^wt/wt^; Lepr^db/db^), RAGE^-/-^ (RAGE^-/-^; Lepr^wt/wt^), and RAGE^-/-^;db/db (RAGE^-/-^; Lepr^db/db^) animals. These animals were allowed to age to 6-8 months of age. Prior to sacrifice, body mass was recorded and blood glucose was measured using blood drawn from the submandibular vein. In, blood was collected from the suborbital vein for HbA1C analysis via an enzymatic assay in a subset of mice (Crystal Chem, Elk Grove Village, IL). The right femur and lumbar vertebrae were dissected following sacrifice, stored at -20C until subsequent analyses.

### 2.2 Small molecule inhibition of RAGE

Inhibition of RAGE were done through the administration of the small molecule antagonist FPS-ZM1.^33,34^ Five month old female db/db mice and wt littermates were used in these experiments. The mice were injected with 1mg/kg of FPS-ZM1 daily for 60 days or the 0.5% DMSO vehicle for 60 days. Four to five mice were used in each of the four groups: vehicle treated wt (0.5% DMSO only), inhibitor treated wt, vehicle treated db/db, and inhibitor treated db/db. Longitudinal measurements of body mass and fasting blood glucose (using blood from tail snips) were taken throughout the duration of the treatment period to examine the effects of FPS-ZM1 on diabetes progression. Additionally, blood was collected from the submandibular vein at the time of sacrifice for HbA1c analysis.

### 2.3 Fluorochrome labeling to evaluate dynamic histomorphometry

Animals were injected with calcine and alizarin 7 and 2 days (RAGE-null) or calcein 2 days (RAGE-inhibition) prior to sacrifice to measure dynamic bone histomorphometry. After sacrifice, the L2 and L3 vertebral bodies were immersed in 70% ethanol (ETOH) after sacrifice and dissection. The vertebral bodies were dehydrated in serial grades of ethanol prior to being embedded in polymethylmethacrylate (PMMA) (Thermo Scientific AAA 130300F, ThermoFisher Scientific). Using a diamond wafer blade (Leica SP1600 Microtome, Leica Microsystems, Inc., Buffalo Grove, IL) two 100um thick transverse cross-sections were taken from approximately 0.2 mm below the cranial growth plate of each vertebral body. Sections were imaged on a confocal microscope (Leica TCS SPEII, 10x lens, 2048x2048, 400Hz, 1.0x zoom). For the RAGE-deficient mice, three channels were imaged: calcein (Ex. 488nm, Em. 502-540nm), alizarin (Ex. 561nm and 633nm, Em. 580-645nm), and bone autofluorescence (Ex. 405, Em. 400-480), while for the RAGE inhibitor-treated mice, two channels were imaged: calcine (495nm ex., 515nm em) and bone autofluorescence (405nm ex, 400-480nm em). Blinded images were analyzed in Bioquant™ (Bioquant Osteo, Nashville, TN) for mineralizing surface per bone surface, the average of two sections was used to calculate mineralizing surface per bone surface (MS/BS) and mineral apposition rate for each animal.

### 2.4 TRAP staining to evaluate osteoclast activity

Both RAGE-deficient mice and inhibitor-treated mice were subject to Tartrate-Resistant Acid Phosphatase (TRAP) staining to quantify osteoclast activity. The L4 vertebral bodies of the RAGE-deficient mice and the L3 vertebral bodies of the inhibitor-treated mice were placed in 4% paraformaldehyde for 24 hours immediately following dissection. Following fixation, bones were decalcified in EDTA for 10 days, serially dehydrated in graded ethanol up to 70%, and embedded in paraffin. Four 10 um thick transverse cross sections were taken from the vertebral bodies, at approximately 1/3 of the vertebral height from the cranial endplate. Sections were imaged on a slide scanner (Hamamatsu NanoZoomer 2.0-HT System, Bridgewater, NJ). Bioquant™ was used to quantify osteoclast surface per bone surface (Oc.S/BS), osteoclast number per bone surface (Oc.N/BS), and osteocyte density for each section. Reported values for each animal are an average of the four sections.

### 2.5 Cortical and Cancellous bone Microstructure by microCT

Both RAGE-deficient mice and inhibitor-treated mice were subjected to microCT analysis and three-point bending to quantify whole-bone mechanics, material properties, and morphology. The L5 vertebral bodies and right femurs were dissected following sacrifice, wrapped in PBS soaked gauze, and stored at -20C. The vertebrae and femurs were thawed for an hour in PBS and imaged in a VivaCT 40 at 10.5um voxel size. Femur and vertebral bone architecture was calculated using a custom MATLAB code.^35^ The cortical bone of the femur mid-diaphysis was analyzed and cortical thickness (Ct.Th), cortical thickness standard deviation (Ct.Th.SD), tissue mineral density (TMD), and polar moment of inertia (pMOI) were all recorded. The cancellous bone was analyzed for bone volume (BV), total volume (TV), bone volume to total volume ratio (BV/TV), connectivity (Conn.), structural model index (SMI), trabecular thickness (Tb.Th.), trabecular thickness standard deviation (Tb.Th.SD), trabecular spacing (Tb.Sp.), trabecular spacing standard deviation (Tb.Sp.SD), and trabecular number (Tb.N).

### 2.6 Whole bone strength mechanical testing of the femora

To evaluate whole-bone mechanical properties, the femurs were subjected to 3-point bending until fracture with a span length of 7.5 mm at a loading rate of 0.1 mm/s (Instron Electropuls, Norwood, MA, USA). Femoral mechanical outcomes include yield load, stiffness, ultimate load, and post yield displacement. Also reported are estimated material properties: Young’s modulus, yield stress, and ultimate stress.^36^

### 2.7 Nanoindentation to evaluate bone matrix nanomechanics

Bones from both RAGE-deficient mice and inhibitor-treated mice were subject to nanoindentation analysis to quantify nano-scale material properties. 2-3 L2 and L3 vertebral bodies from these groups (N=20) were embedded in PMMA. The PMMA-embedded blocks were ground to expose the transverse cross-section approximately 0.4 mm from the cranial growth plate using silicon carbide sheets. The face of the block was then polished to a 0.05 um finish using increasing grits of silicon carbide sheets and 1 um and 0.05 um alumina suspensions on a polishing wheel. A Hysitron 980 nanoindenter with a Berkovich probe was used for testing (Bruker-Hysitron, Eden Prairie, MN USA). A trapezoidal loading curve was utilized beginning with a 800uN/s loading rate until the maximum load of 10mN was reached, the maximum load was maintained for 30s, followed by an unloading rate of 800uN/s. For the second loading condition a trapezoidal loading curve was loaded at a loading rate of 80uN/s until a maximum load of 1mN was reached, maximum load was maintained for 30s, followed by an unloading rate of 80uN/s. Fifteen indentations were made in each sample for each loading condition. Indents were randomly distributed across the cancellous boney regions with a minimum distance of 10 um between indents. To clean the probe a polished piece of aluminum was indented three times after every two samples. The machine was recalibrated halfway through testing using fused quartz to account for effects of tip dulling.

To calculate material properties from the nanoindentation data two methods were utilized. First, the Oliver-Pharr method was used to calculate the reduced modulus (𝐸_𝑅_, Eq 1) and contact hardness (Hc, Eq 2). In the case of bone reduced modulus is equivalent to indentation modulus (E’) and is calculated based on the unloading stiffness (S) and contact area (𝐴_𝑐_). Contact hardness is the peak load (P_max_) divided by the contact area (𝐴_𝑐_).^37^

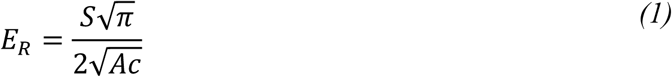

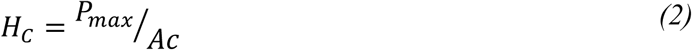

Since AGEs are known to crosslink the organic matrix, and the organic matrix is primarily responsible for the viscoelastic behavior of bone, we also measured the viscoelastic behavior of the bone matrix. The 30s dwell during the load function allows for analysis of time-dependent mechanical properties of the bone. To analyze this data we utilized the viscoelastic-plastic analysis (VEP) to determine the viscous response during the holding period. To calculate the indentation viscosity (𝜂_Q_) the displacement-time curve during the holding period (ℎ^𝑐𝑟𝑒𝑒𝑝^(𝑡)) was fit based on equation 3. The calculation assumes a linear creep rate, for a more accurate analysis only the last 5/6ths of the hold period was fit, the beginning of which is defined as 𝑡_1_. Variables in the equation include the loading rate (k), time until maximum load (𝑡_𝑟_), displacement at 𝑡_1_(ℎ(𝑡_1_)), and a geometric constant for a perfect Berkovich tip (𝛼_3_). This data was fit in MATLAB using a custom MATLAB code (S1).^37,38^

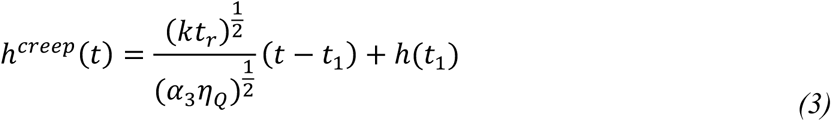

### 2.8 Fluorometric assay to measure the accumulation of AGEs in the bone matrix

Bone from RAGE-deficient mice and RAGE-inhibitor-treated mice were subject to the total fluorometric AGE assay to quantify density of AGE crosslinks in bone collagen. The L5 vertebral bodies and right femurs were used, after removal of the facet joints from the vertebrae and isolation of a 2mm section of the diaphysis adjacent to the femur break site. For both vertebrae and femur samples, the following procedure was used. Samples were decalcified in Immunocal™ for 3 days, hydrolyzed in 6N HCl at 65°C for 3 hours, and then samples were pipetted into a 96 well plate in triplicate and HCl was allowed to evaporate overnight. Samples were resuspended in PBS for an hour and then imaged with a BioTek Synergy plate reader at emission wavelength 370nm with a capture wavelength of 440nm and standardized to quinine standards with known concentrations. These data were normalized to the amount of collagen in the bone determined by the amount of hydroxyproline. Hydroxyproline was quantified using a chloramine-T colorimetric assay standardized to commercially available hydroxyproline standards (BioTek Synergy, absorbance 560 nm).^39^

### 2.9 High Resolution X-ray Microscopy of Osteocyte Lacunae

Lacunar morphology was assessed only in RAGE-deficient mice. This choice was based on the premise that changes in the lacunar microenvironment manifest over longer time scales, which are more appropriately captured in the RAGE⁻/⁻ model. The L2 vertebral body was dissected and stored at -20°C in PBS soaked gauze in a subset of the mice (n = 3-4/group) for XRM. Facet joints and vertebral endplates were resected, and the vertebral column was put into an 2mm diameter holder with PBS and topped with wax to keep the bone hydrated during imaging. XRM imaging was performed by using a 4x scout scan to identify a 0.067mm^3^ ROI of cancellous bone in the middle of the vertebral body, 500 nm below the cranial endplate (Figure 4A). The region was imaged at 455.8 nm resolution using a 20x objective (no filter, bin of 2, 4keV, 3W). Image analysis was performed in Dragonfly ORS (Montreal, Quebec). Lacunae were identified as low attenuating voxels within the bone area with volumes greater than 5*10^-8^ mm^3^. Lacunae and partially embedded lacunae were identified and analyzed separately as functions of bone volume (BV) and bone surface area (SA) respectively (Figure 4C-I). Size and shape of the lacunae were also analyzed; reported here are lacunar volume (Lac.V), lacunar volume per surface area (Lac.V/SA), aspect ratio, and min orthogonal to max feret diameter.

### 2.10 Time-lapsed microCT to evaluate 3D dynamic histomorphometry

The RAGE inhibitor-treated mice were subject to longitudinal microCT analysis in order to quantify serial bone changes over the 2-month treatment period with FPS-ZM1. The L5 vertebral bodies were imaged with a VivaCT40 (Scanco medical AG, VivaCT40, Bassersdorf, Switzerland) at 10.5 um voxel size, mice were anesthetized with isofluorane for approximately 20 minutes for each scan. DICOM stacks were exported from the Scanco software for cancellous bone isolation in MATLAB using a custom GUI^35^ and then analysis in Dragonfly ORS (Version 2024.1 for Windows, Comet Technologies Canada Inc., Montreal, Canada). A number of the scans had to be excluded due to motion artifacts, resulting in n=3 mice/group that had a full set of microCT images for analysis. The cancellous bone area of the vertebral column was isolated to a region approximately 40um away from the growth plates. The isolated cancellous bone DICOM stacks were uploaded to Dragonfly ORS for image registration and analysis. The images were registered in 3-D space using the image registration plugin in Dragonfly ORS, the mutual information matching process was used to match the week 4 and week 8 DICOM stacks to the week 0 DICOM stack. The images were registered with a smallest step size of 0.005 um in each direction and 0.05 degrees for rotation. Based on the image registration the volume of interest (total volume) was further reduced to only the volume that was consistent throughout all imaging timepoints, the range in total volume analyzed was between 1.01mm^3^ to 2.13 mm^3^, average of 1.74 mm^3^. This was necessary as it was not possible to image the entire vertebral body in the short scan time, so only overlapping regions were able to be analyzed. A bone mask was defined for each imaging timepoint based on a consistent threshold measured in hounsfield units, the bone masks were then smoothed with a kernel size of 5 to reduce partial volume effects. Bone volume to total volume ratio of the region of interest was reported at 0 weeks, 4 weeks after the start of treatment, and 8 weeks after the start of treatment. To further characterize the changes in BV/TV the mineralized bone volume divided by total bone volume, eroded volume divided by total bone volume, steady-state bone volume divided by total bone volume were reported for the change from week 0 to week 4 and the change from week 4 to week 8.

### 2.11 Statistics

Statistical analyses were performed in GraphPad PRISM 10 (GraphPad Software, Boston, Massachusetts USA). Terminal measurements, excluding nanoindentation tests, were analyzed with a two-way ANOVA testing for effects for testing for effects of diabetes and RAGE deletion or inhibition (treatment). Longitudinal data was analyzed using a repeated measures three-way ANOVA testing for effects of diabetes, RAGE deletion or inhibition (treatment), and time (length of treatment).

Statistical analyses for the nanoindentation measurements were performed in MATLAB after removal of statistical outliers. Statistical outliers were removed using the ROUT method in GraphPad Prism with a 1% Q, outlier tests were run individually for each sample. A linear mixed effects model was utilized to be able to account for random effects of mouse on the data set.^40,41^ Diabetes and RAGE^-/-^ treatment were both treated as fixed effects in the model whereas mouse was treated as a random effect, the model was run on each outcome individually after normalization by subtracting the average of all of the data points.^42^ When main effects or interaction effects had a p-value <0.10 post-hoc analyses were also run as a linear mixed model in MATLAB keeping each mouse as a random effect.

## 3. Results

### 3.1 The ablation of RAGE does not affect the metabolic phenotypes of diabetes but prevents the accumulation of AGEs in bone

Leptin receptor-deficient mice developed obesity, hyperglycemia, and elevated hemoglobin A1c (HbA1c), consistent with the canonical diabetic phenotype of db/db animals.^30^ The lack of RAGE signaling in these animals did not affect the penetrance of the diabetes phenotype (Figure 1A-C). However, RAGE-deficient diabetic animals were protected from the accumulation of AGEs in their bone matrix (Figure 1D). The ablation of RAGE prevented the diabetes-mediated accumulation of AGEs in RAGE^-/-^;db/db animals. In the RAGE inhibition experiments, diabetes was associated with increased body mass, blood glucose, HbA1c, and vertebral AGE concentration (Figure 1E-G). FPS-ZM1 treatment did not affect AGE concentrations in the vertebra (Figure 1H). Longitudinal measurements in the RAGE inhibition experiments suggest that FPS-ZM1 treatment may mitigate the diabetic increase in blood glucose over time (Figure S2).

**Figure 1.**
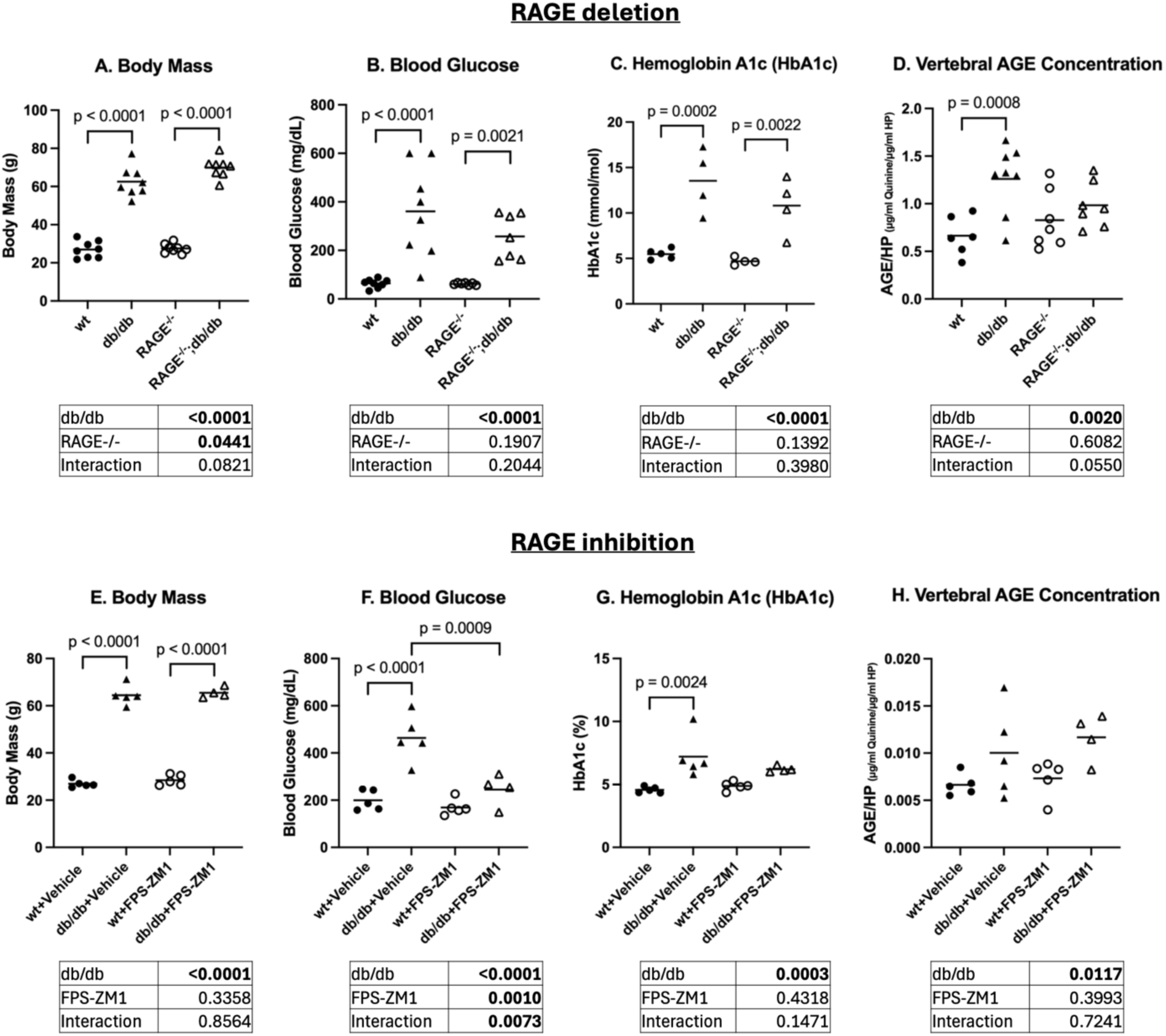
Leptin receptor-deficient mice exhibit clinical metrics of diabetes which are mitigated by RAGE inhibition, while RAGE deletion prevents vertebral AGE accumulation in db/db animals. Increased body mass (A), blood glucose (B), and HbA1c (C) in diabetic animals compared to respective controls. (D) Vertebral AGE concentration was elevated in db/db animals compared to wt but not in RAGE^-/-^;db/db compared to RAGE^-/-^. Increased body mass (E) in diabetic animals compared to respective controls. Blood glucose (F) and HbA1c (G) are elevated with diabetes in vehicle-treated mice but not inhibitor-treated mice. (H) Diabetes is associated with increased vertebral AGE concentration across all groups with no effect of RAGE inhibition.

### 3.2 Loss of RAGE in diabetic mice improves trabecular microarchitecture

Animals with diabetes have higher bone volume fraction than respective controls (Figure 2A, B). This increase in BV/TV is characterized by an increase in trabecular connectivity and a decrease in trabecular spacing with no change in trabecular thickness (Figures 2C-E). As a consequence of preventing bone accrual in the diabetic animals, the deletion of RAGE also concomitantly decrease trabecular connectivity, which has been previously reported in mouse femoral trabecular bone while maintaining the trabecular thickness (Figure 2B-D).^43^ In the cortical shell of the vertebra, diabetes negatively affects TMD but not mean thickness while RAGE deletion has no effect on either cortical parameter (Figure 2G).

**Figure 2.**
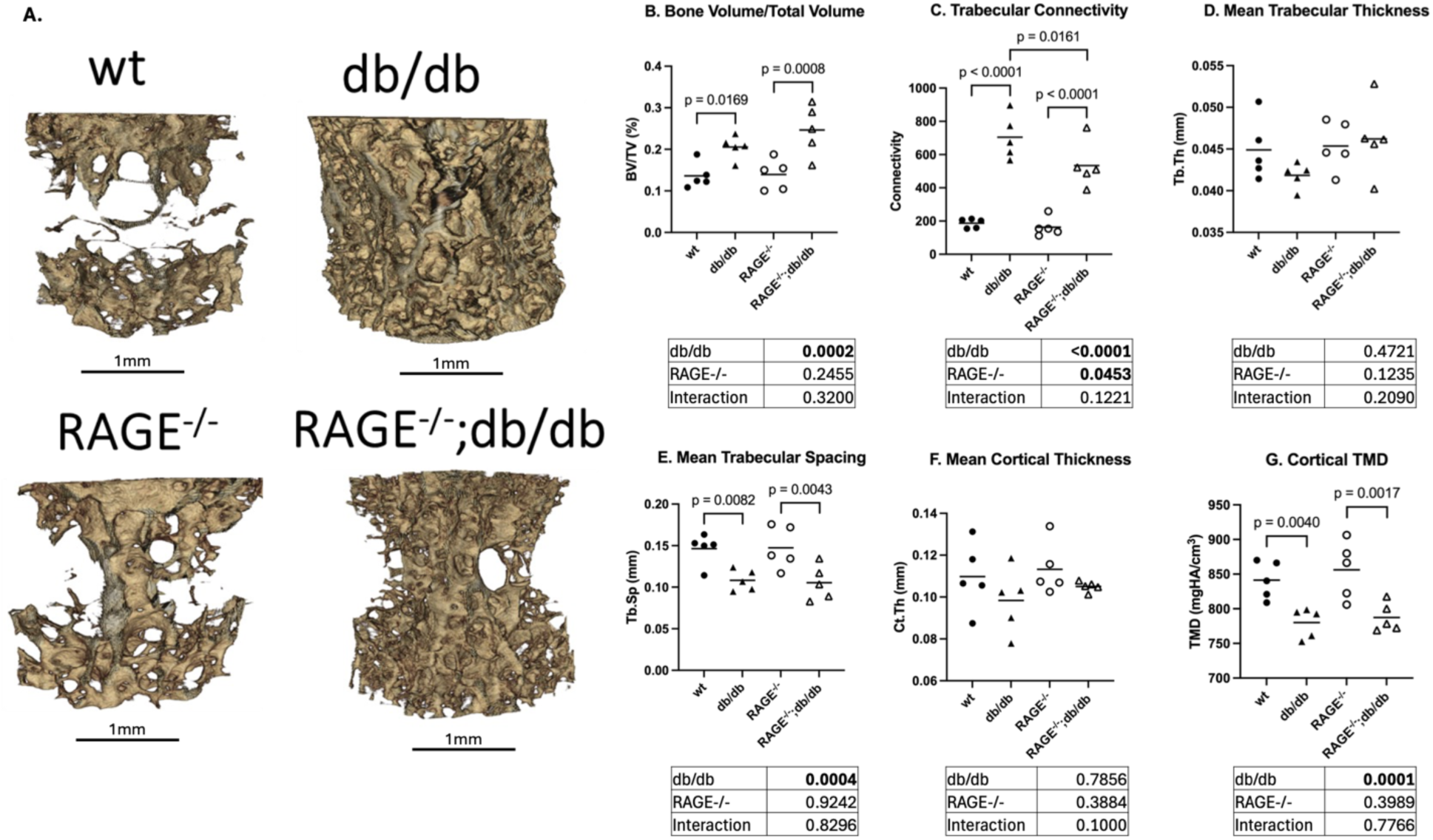
RAGE deletion modifies aspects of trabecular microarchitecture but not cortical bone thickness or TMD. (A) Representative 3D renderings from each group. Increased BV/TV (B) and connectivity (C) in diabetic animals, no change in trabecular thickness (D) and mean cortical shell thickness (F) with diabetes, and decreased trabecular spacing (E) and cortical shell TMD associated with diabetes (G).

### 3.3 Loss of RAGE is associated with increased osteoblast activity and, in diabetic animals, may improve osteoclast function

Across all groups, diabetes was associated with decreased MS/BS and RAGE deletion was associated with increased MS/BS (Figure 3B). RAGE deletion was associated with increased MS/BS across all groups. Osteoclast activity was not significantly affected by diabetes in this sample, although most previous reports find inhibited osteoclast activity in diabetic animals^9,10^. However, despite lacking significance, Oc.S/BS may be affected by RAGE deletion in a diabetes-dependent manner (p=0.0970). RAGE deletion in wild type animals resulted in decreased Oc.S/BS, while in diabetic animals RAGE deletion resulted in increased Oc.S/BS. This divergence results in the RAGE^-/-^;db/db group trending higher in Oc.S/BS compared to the db/db group (p=0.0873).

**Figure 3.**
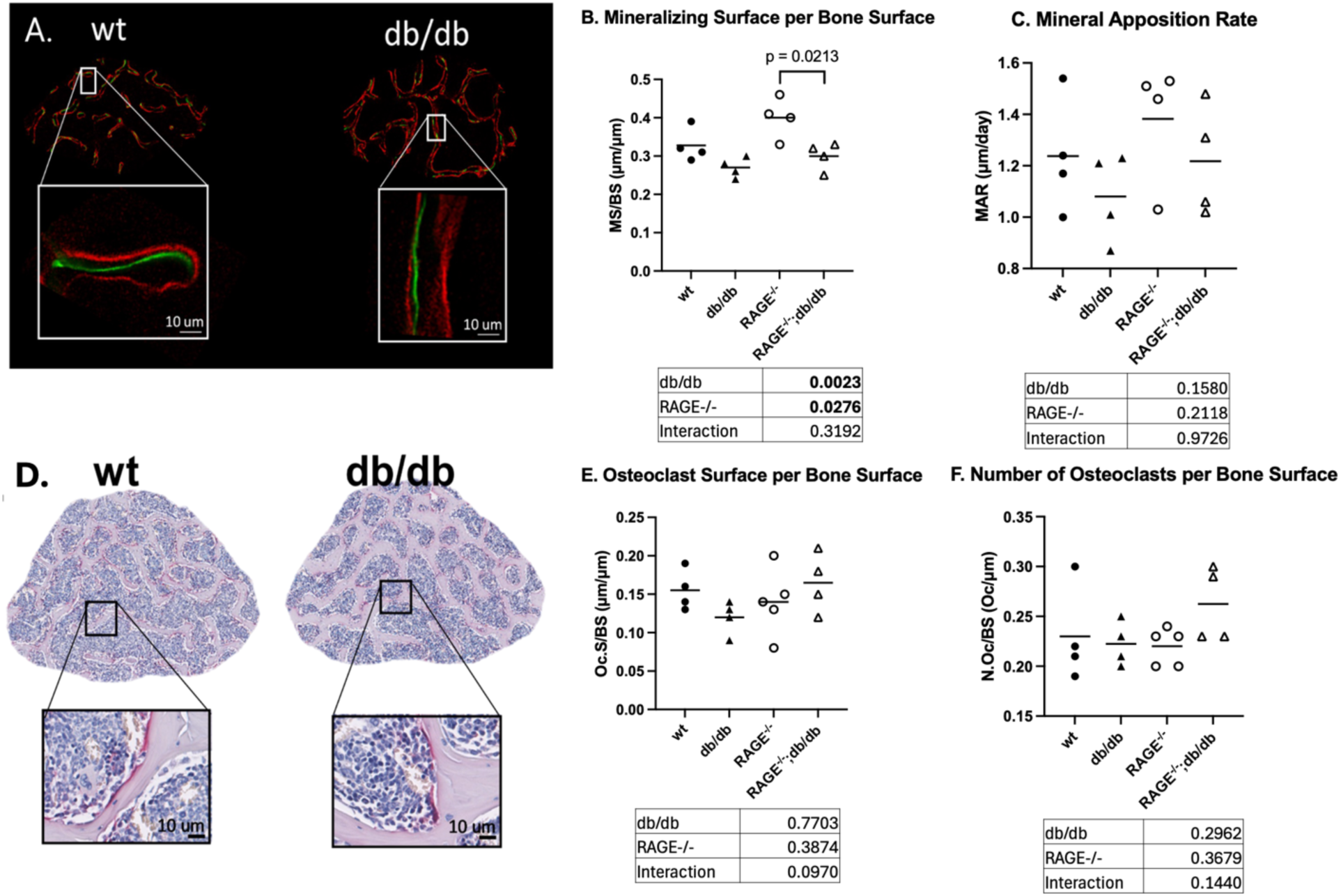
Diabetes and RAGE deletion have opposing effects on osteoblast activity, but neither have significant effects on osteoclast activity. Representative images of wt and db/db (A) calcine and alizeran labeling and (D) TRAP counterstained with H&E. Bone formation measures showed (B) increased MS/BS in RAGE-deficient animals overall and (C) no change in MAR. Osteoclast activity measures showed no significant changes in Oc.S/BS (E) or osteoclast number (F).

### 3.4 Loss of RAGE improves osteocyte density and lacunar morphology in diabetic animals

The db/db animals with intact RAGE signaling had diminished number of osteocyte lacunae per bone volume compared to wt, while there was no significant difference between RAGE-deficient groups (Figure 4G). RAGE deletion was associated with increased lacunar volume across all groups, with the RAGE^-/-^ and RAGE^-/-^;db/db groups exhibiting significantly larger lacunar volume than the respective controls. No significant effects of diabetes or treatment were observed in lacunar volume to surface ratio, although the increased mean lacunar volume to surface ratio in db/db mice compared to wt mice was trending toward significance (p=0.080) with no difference between the RAGE-deficient groups. Across all groups, diabetes was associated with decreased lacunar aspect ratio and RAGE deletion was associated with an increase. However, RAGE deletion appears to increase aspect ratio only in wild-type mice. Min orthogonal to max feret diameter is another measure of lacunar shape which describes the elongation of a particle. Min ortho/max feret was increased in the db/db group compared to wt, indicating less elongated lacunae. No significant difference in min ortho/max feret was observed between the RAGE- deficient groups. In terms of diabetic osteocyte morphology, RAGE deletion improves lacunar min ortho/max feret and may improve lacunar volume to surface ratio. RAGE deletion also had effects on lacunar volume that were independent of diabetic status and on lacunar aspect ratio that were limited to wild-type animals.

### 3.5 RAGE inhibition mitigates bone accrual in diabetic animals

In the RAGE inhibitor-treated mice, the detrimental effects of diabetes on vertebral trabecular microarchitecture were consistent with the deficiencies observed in RAGE-deficient mice. RAGE inhibition protected against elevated bone volume fraction due to diabetes (Figure 5B), which was not observed in the RAGE deletion study. RAGE inhibition did not alter trabecular connectivity in the diabetic animals, in contrast to the effects of RAGE deletion (Figure 5C). Diabetes impaired cortical shell TMD with no effect of FPS-ZM1, which aligns with our findings in the RAGE-deficient mice (Figure 5G). To further investigate the beneficial effect of RAGE inhibition on BV/TV, we next analyze the longitudinal measurements taken throughout the duration of FPS-ZM1 treatment.

**Figure 4.**
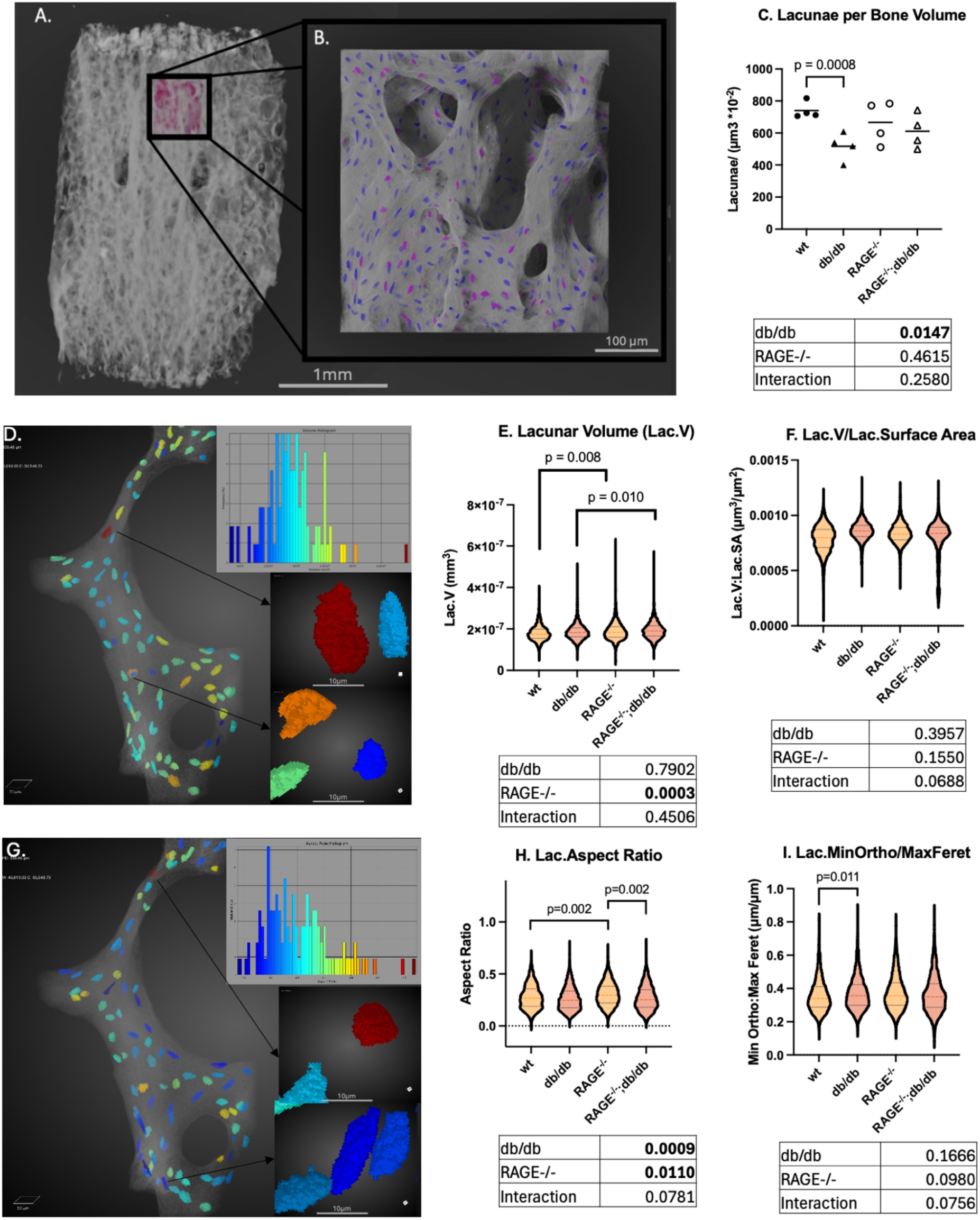
Diabetes modifies osteocyte density and lacunae shape, with RAGE deletion protecting against both impairments. Representative region on interest (A) 4x scout scan, (B) high resolution scan with lacunae highlighted in blue and surface lacunae in pink. (C) Reduced lacunar density in cancellous bone of db/db animals. (D) Representative region depicting difference between smaller and larger lacunae. (E) Lacunar volume is increased in RAGE^-/-^ animals compared to wt, and RAGE^-/-^;db/db compared to db/db. (F) Trending increase in lacunar volume to surface area ratio in db/db animals compared to wt. (G) Representative region depicting high to low aspect ratio lacunae. (H) Lacunar aspect ratio is increased in RAGE^-/-^ and RAGE^-/-^;db/db animals. (I) Lacunar min ortho diameter to max feret diameter is decreased in db/db animals compared to wt and increased in RAGE^-/-^.

**Figure 5.**
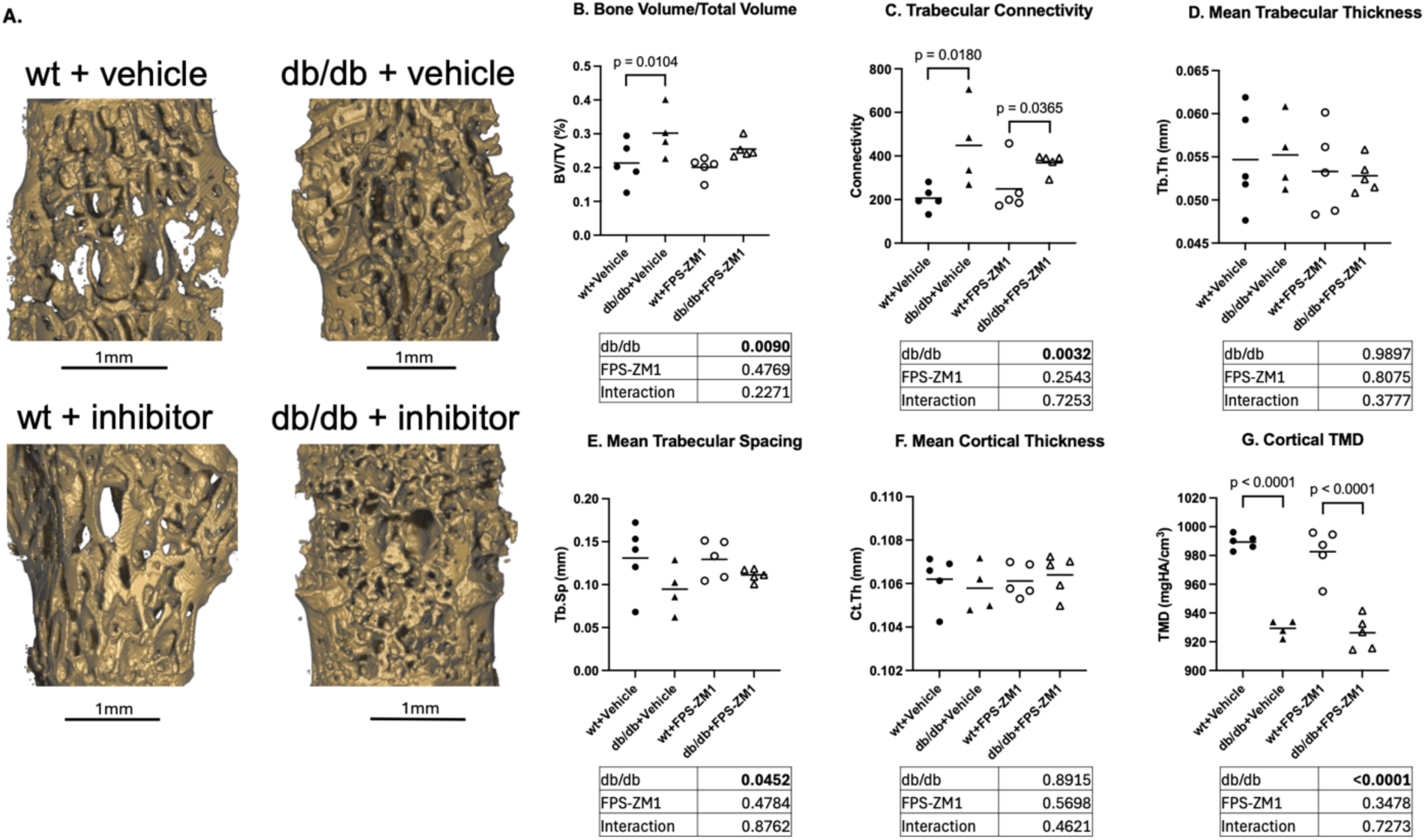
RAGE inhibition improves diabetic bone volume fraction but does not improve other aspects of trabecular microarchitecture. (A) Representative 3D renderings from each group. Increased vertebral BV/TV (B) and connectivity (C) in diabetic animals, no change in trabecular thickness (D) and mean cortical shell thickness (F), and decreased trabecular spacing (E) and cortical shell TMD (G).

### 3.6 RAGE inhibition restores osteoclast activity and osteocyte density in diabetic animals

Time-lapse MicroCT showed that the db/db group gained a significant amount of bone volume fraction between four to five months of age (4-8 weeks after the start of FPS-ZM1 treatment, Figure 6A). This time-dependent increase in BV/TV was not present in the FPS-ZM1- treated diabetic animals. Between four to five months of age, the db/db group experienced a significant decrease in erosion percentage compared to the 3-4-month time point, which temporally aligns with the elevation in BV/TV (Figure 6D). This resulted in an increase in steady-state bone, or bone that has not been eroded, for the corresponding time point in the db/db group (Figure 6E).

**Figure 6.**
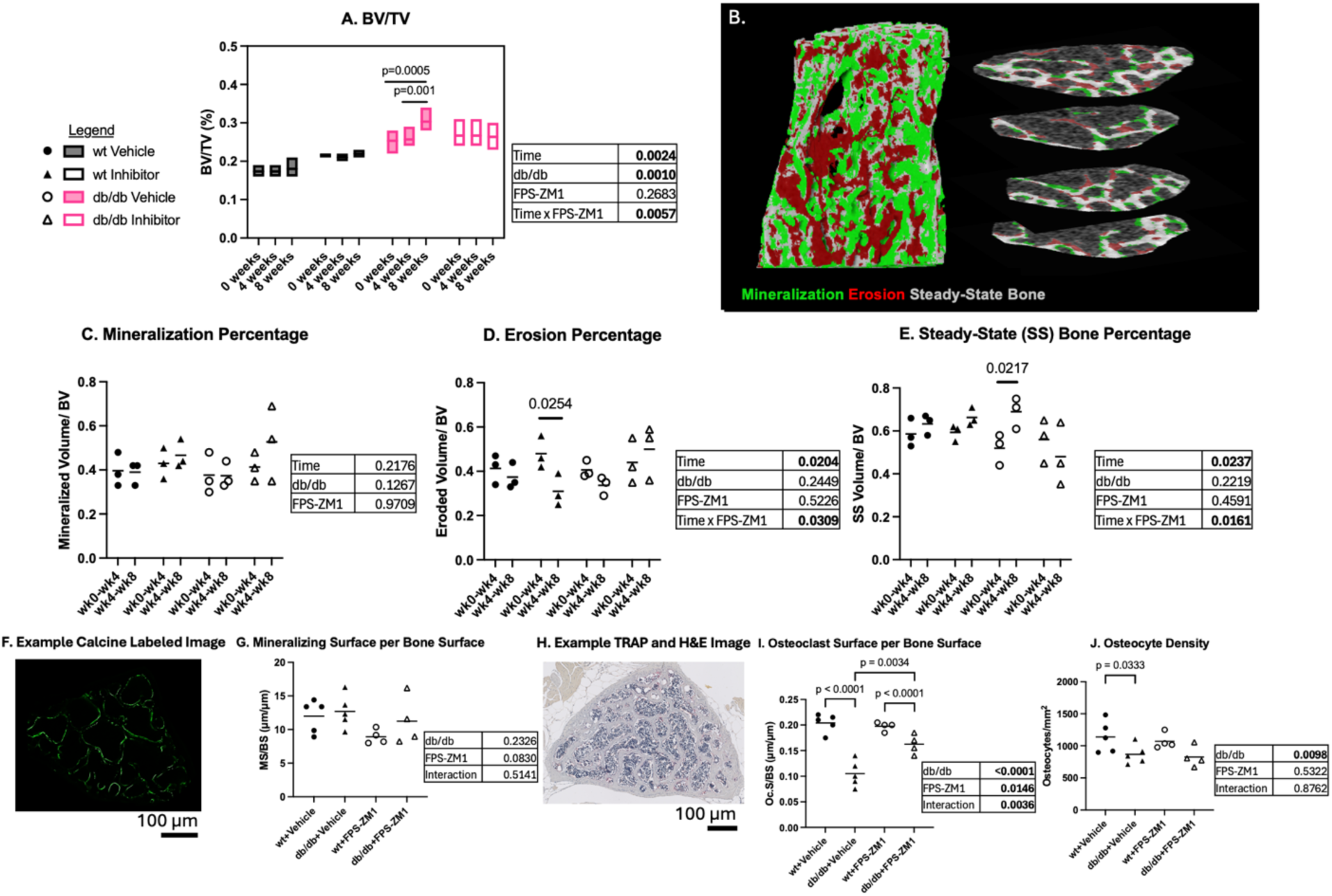
RAGE inhibition in diabetic mice alters bone volume fraction in a time-dependent manner and increases erosion percentage and osteoclast activity. (A) Bone volume to total volume ratio was increased due to diabetes overall, with a marked increase over time in the db/db vehicle group. (B) Example rendering showing newly mineralized bone, newly eroded areas, and steady-state bone between week 0 and week 4 of treatment. (C) Mineralization volume normalized to initial bone volume was not changed due to diabetes, time, or treatment. (D) Erosion volume normalized to initial bone volume was decreased over time in the db/db vehicle group. (E) Steady-state bone percentage was increased over time in the db/db vehicle group. (F) Example Calcine labeled cross-section and (G) mineralizing surface to bone surface (MS/BS) showed no changes due to treatment or diabetes. (H) Example TRAP and calcine histological cross-section, and (I) osteoclast surface to bone surface (Oc.S/BS) was reduced in the db/db vehicle group compared to the db/db inhibitor treatment group. (J) Reduced osteocyte density was noted in db/db animals vs wild type with no effect of treatment.

Diabetes was not associated with changes in osteoblast activity, but it was associated with reduced osteoclast activity and osteocyte density across all groups. Diabetic animals treated with FPS-ZM1 exhibited increased osteoclast activity compared to vehicle-treated controls (Figure 6K). There was no statistical difference in osteocyte activity between the FPS-ZM1 treated groups, while the db/db+vehicle group had reduced osteocyte density compared to the wt+vehicle group (Figure 6L).

### 3.7 RAGE deletion and inhibition improve bone viscoelastic properties in diabetic animals

Diabetes was associated with increased indentation modulus in the RAGE-deficient mice across all groups, with higher indentation modulus in the db/db group compared to wt (Figure 7E). In contrast to the groups with intact RAGE signaling, there was no statistical difference in indentation modulus between RAGE-deficient groups. Contact hardness was increased in the RAGE^-/-^;db/db group compared to db/db and RAGE^-/-^ groups (Figure 7F). RAGE inhibition yielded the opposite outcome: contact hardness was decreased in the db/db+FPS-ZM1 group compared to db/db+vehicle and wt+FPS-ZM1 groups (Figure 7I). Indentation viscosity was increased in the RAGE^-/-^;db/db group compared to the db/db group, and the significant interaction value indicates that RAGE deletion reversed the detrimental effect of diabetes on indentation viscosity (Figure 7G). RAGE inhibition yielded a similar outcome: indentation viscosity was increased in the db/db+FPS-ZM1 group compared to the db/db+vehicle group, again with a significant interaction effect (Figure 7J).

**Figure 7.**
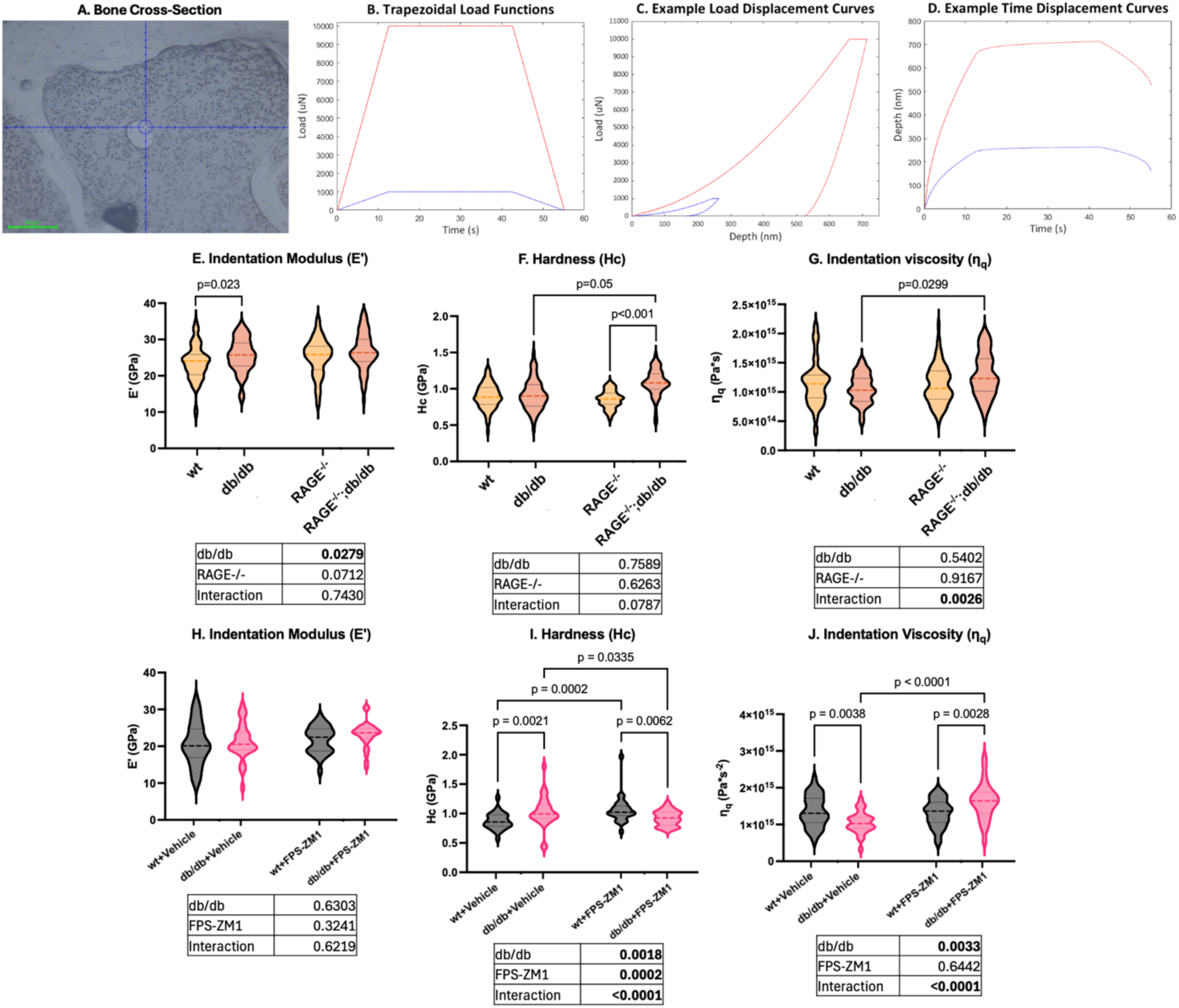
Nanomechanical impairments of the bone matrix mediated by diabetes are prevented by RAGE deletion and remediated by RAGE inhibition, with both improving bone nanomechanical behavior. (A) Sample cross-section used to identify indentation location. (B) Sample trapezoidal load function (B), load displacement curve (C), time displacement curve (D) with 10mN indent. (E) Indentation modulus 10mN for RAGE deletion, effect of diabetes with db/db higher than wt. (F) Hardness for RAGE deletion, RAGE^-/-^;db/db higher than db/db and RAGE^-/-^. (G) Indentation viscosity for RAGE deletion, significant interaction effect, RAGE^-/-^;db/db higher than db/db. (H) Indentation modulus for RAGE inhibition, no significant outcomes. (I) Hardness for RAGE inhibition, effect of diabetes, treatment, and interaction, db/db+vehicle higher than wt+vehicle, db/db+FPS-ZM1 lower than db/db+vehicle and wt+FPS-ZM1. (J) Indentation viscosity for RAGE inhibition, significant diabetes and interaction effects with db/db+vehicle lower than wt+vehicle and db/db+FPS-ZM1 higher than db/db+vehicle and wt+FPS-ZM1.

### 3.8 RAGE deletion prevents pathological changes in femoral bone mechanics, but RAGE inhibition does not fully rescue the diabetes mediated changes

Diabetic control femurs were mechanically inferior as indicated by several mechanical outcomes. The deletion of RAGE in diabetic animals prevented deteriorations in stiffness, yield load, ultimate load, work to fracture, and yield stress (Table 1). RAGE deletion also prevented a decrease in cortical thickness in diabetic animals, but RAGE deletion did not affect the AGE concentrations in the femur. RAGE inhibition rescued yield load but no other mechanical paramaters (Table 2). RAGE inhibition restored the cortical thickness and pMOI and reversed the AGE accumulation in the femur.

**Table 1.** RAGE deletion prevents diabetes-associated decline in multiple cortical bone mechanical parameters and cortical thickness. Mechanical improvements to stiffness, yield load, ultimate load, and work to fracture were noted in RAGE-null diabetic animals. For each group and outcome the mean ± standard deviation is reported, and the outcomes of statistical comparisons are also denoted in the table. P-values less than or equal to 0.05 are bolded and significant post-hoc differences between groups are denoted with matching letters (a or b).

|  | Group Mean |  |  |  | ANOVA Effects |  |  |
| --- | --- | --- | --- | --- | --- | --- | --- |
| Mechanics | wt | db/db | RAGE-/- | RAGE-/-;db/db | Diabetes | RAGE-/- | Interaction |
| Stiffness (N/mm) | 118.31 ± 33.87 **a** | 89.34 ± 10.68 **a** | 108.99 ± 12.57 | 103.95 ± 20.88 | **0.0436** | 0.7424 | 0.1466 |
| Yield Load (N) | 13.13 ± 4.25 **a** | 9.49 ± 1.47 **a** | 11.01 ± 1.24 | 11.60 ± 2.24 | 0.1232 | 0.9978 | **0.0370** |
| Ultimate Load (N) | 19.52 ± 3.61 **a** | 14.51 ± 2.22 **a** | 16.97 ± 1.27 | 16.86 ± 2.16 | **0.0106** | 0.9141 | **0.0137** |
| Post Yield Displacement (mm) | 0.43 ± 0.16 | 0.31 ± 0.17 | 0.49 ± 0.16 | 0.51 ± 0.24 | 0.4609 | 0.0872 | 0.3474 |
| Work to Fracture (N*mm) | 6.30 ± 1.75 **a** | 3.55 ± 2.18 **a,b** | 6.79 ± 2.00 | 6.50 ± 1.84 **b** | 0.0589 | **0.0341** | 0.1200 |
| Young's Modulus (N/mm<sup>2</sup>) | 8232 ± 2493 **a** | 6230 ± 1244 **a** | 9516 ± 653 **b** | 7178 ± 906 **b** | **<0.0001** | 0.1120 | 0.4543 |
| Yield Stress (N/mm<sup>2</sup>) | 123.05 ± 38.48 | 94.40 ± 19.81 | 123.39 ± 12.51 | 111.08 ± 29.08 | 0.0505 | 0.4003 | 0.4188 |
| Ultimate Stress (N/mm<sup>2</sup>) | 183.08 ± 27.22 **a** | 142.43 ± 14.18 **a** | 190.67 ± 18.62 **b** | 158.98 ± 18.35 **b** | **<0.0001** | 0.1296 | 0.5649 |
|  | Group Mean |  |  |  | ANOVA Effects |  |  |
| Bone Quality | wt | db/db | RAGE-/- | RAGE-/-;db/db | Diabetes | RAGE-/- | Interaction |
| Cortical Thickness (mm) | 0.2372 ± 0.02 **a** | 0.1893 ± 0.02 **a** | 0.2197 ± 0.02 | 0.2088 ± 0.01 | **<0.0001** | 0.5837 | **0.0229** |
| pMOI | 0.3694 ± 0.07 **a** | 0.3623 ± 0.07 | 0.2882 ± 0.03 **a,b** | 0.3620 ± 0.06 **b** | 0.1382 | 0.0728 | 0.0753 |
| AGEs (ug/ml)*10<sup>2</sup> | 0.4239 ± 0.21 | 0.6989 ± 0.45 | 0.4366 ± 0.12 | 0.3877 ± 0.11 | 0.3218 | 0.1947 | 0.1609 |

**Table 2.** RAGE inhibition improved yield load and femur AGE concentration in diabetic animals. Diabetic animals treated with FPS-ZM1 exhibited decreased cortical thickness and pMOI compared to wt+FPS-ZM1 animals. For each group and outcome the mean ± standard deviation is reported, and the outcomes of statistical comparisons are also denoted in the table. P-values less than or equal to 0.05 are bolded and significant post-hoc differences between groups are denoted with matching letters (a or b).

|  | Group Mean |  |  |  | ANOVA Effects |  |  |
| --- | --- | --- | --- | --- | --- | --- | --- |
| Mechanics | wt Vehicle | db/db Vehicle | wt Inhibitor | db/db Inhibitor | Diabetes | Treatment | Interaction |
| Stiffness (N/mm) | 119.6 ± 22.9 **a** | 98.5 ± 3.1 | 142.4 ± 15.3 **a,b** | 91.3 ± 8.2 **b** | **0.0002** | 0.3039 | 0.0576 |
| Yield Load (N) | 17.1 ± 3.7 **a** | 12.7 ± 0.9 **a** | 16.4 ± 3.2 | 12.6 ± 1.1 | **0.0063** | 0.7400 | 0.7993 |
| Ultimate Load (N) | 20.2 ± 2.4 **a** | 15.3 ± 1.0 **a** | 22.3 ± 1.4 **b** | 15.4 ± 1.9 **b** | **<0.0001** | 0.2127 | 0.2720 |
| Post Yield Displacement (mm) | 0.50 ± 0.20 | 0.50 ± 0.20 | 0.42 ± 0.07 | 0.57 ± 0.18 | 0.3505 | 0.9503 | 0.3188 |
| Work to Fracture (N*mm) | 9.26 ± 3.68 | 6.38 ± 2.11 | 7.67 ± 1.94 | 7.56 ± 2.27 | 0.2541 | 0.8713 | 0.2905 |
| Young's Modulus (N/mm<sup>2</sup>) | 6737 ± 1364 | 5796 ± 4266 | 7275 ± 1001 **a** | 4954 ± 224 **a** | **0.0024** | 0.7359 | 0.1407 |
| Yield Stress (N/mm<sup>2</sup>) | 137.7 ± 29.4 | 112.4 ± 4.6 | 120.7 ± 27.6 | 99.7 ± 5.9 | **0.0426** | 0.1735 | 0.8424 |
| Ultimate Stress (N/mm<sup>2</sup>) | 162.5 ± 18.2 **a** | 135.3 ± 1.6 **a** | 163.6 ± 10.1 **b** | 121.9 ± 10.7 **b** | **<0.0001** | 0.3036 | 0.2309 |
|  | Group Mean |  |  |  | ANOVA Effects |  |  |

Table 2. RAGE inhibition improved yield load and femur AGE concentration in diabetic animals.
| Bone Quality | wt Vehicle | db/db Vehicle | wt Inhibitor | db/db Inhibitor | Diabetes | Treatment | Interaction |
| --- | --- | --- | --- | --- | --- | --- | --- |
| Cortical Thickness (mm) | 0.22 ± 0.03 | 0.20 ± 0.02 | 0.24 ± 0.01 **a** | 0.19 ± 0.01 **a** | **0.0009** | 0.5381 | 0.1319 |
| pMOI | 0.45 ± 0.05 | 0.43 ± 0.04 | 0.51 ± 0.02 **a** | 0.438 ± 0.071 **a** | **0.0384** | 0.1295 | 0.3181 |
| AGEs (ug/ml)*10<sup>2</sup> | 0.4159 ± 0.07 **a** | 0.6825 ± 0.18 **b** | 0.5439 ± 0.07 **a** | 0.4181 ± 0.07 **b** | 0.1951 | 0.2073 | **0.0043** |

## 4. Discussion

The results of this study showed a critical role of RAGE signaling in the detrimental effects of T2D on bone properties. These findings extend our understanding of the role of RAGE as a regulator of bone cell homeostasis and clarify the role of RAGE in diabetic bone quality.

AGE formation is accelerated in T2D due to hyperglycemia, as the abundance of sugars bind to the amino acid residues in proteins such as bone. AGEs impair bone quality through both direct crosslinking on the bone matrix and effects on bone cells. The data here suggests that RAGE deletion and inhibition can mitigate the increases in collagen glycation due to T2D. This aligns with previous reports indicating that RAGE activation can perpetuate AGE accumulation by downregulating protective mechanisms such as glyoxalase 1, which normally clear precursors of AGE formation.^45^ This improved bone composition may help account for the beneficial effects of RAGE deletion and inhibition on nanoindentation outcomes in diabetic animals. Prior research has found that AGE concentration is inversely related to bone post-yield mechanics at the macroscale.^45^ At the micro-scale, the τ_2_ relaxation time from nanoindentation correlated with decreased pentosidine levels in the bone in a chronic kidney disease model.^46^ Our findings complement the existing literature by showing how AGE concentration can be reduced by targeting RAGE in diabetic animals, which may contribute to the improved matrix and whole-bone mechanics we observed in this study.

Several studies have investigated the effect of directly targeting AGE crosslinking on bone mechanical properties. The aminoguanidine has shown efficacy ex vivo in reducing AGE density in cortical bone and improving bone mechanical properties.^47^ AGE cross-link breakers have also shown to be effective in vitro for improving bone mechanics and AGE concentration.^48,49^ Few studies have investigated the effect of reducing AGE concentration on bone mechanical properties in vivo. One study using a mouse model of chronic kidney disease showed that treatment with an AGE cross-link breaker reduced fAGE concentration with no change in bone mechanical properties.^50^ In this study, we demonstrated in a mouse model of T2D that in vivo deletion and inhibition of RAGE are viable approaches to address matrix AGE accumulation and bone biomechanics. However, the relative contributions of AGE crosslinking and bone cell dysfunction to the impaired bone quality associated with T2D remain unclear.

To investigate the cellular side of the AGE-RAGE problem, we employed a high-resolution approach for analyzing bone cell behavior that resulted in numerous novel findings. Our results show that T2D impairs the activity of all bone cells, and that RAGE deletion and inhibition confer benefits primarily to osteoclasts and osteocytes. In diabetic animals, RAGE deletion appears to improve osteocyte density and lacunar morphology, as well as osteoclast activity. This modified osteocyte and osteoclast function may explain the decreased trabecular connectivity observed in the RAGE^-/-^;db/db animals. Prior research has demonstrated an inverse relationship between connectivity and stiffness in trabecular bone^51^, which is consistent with our findings of improved whole-bone mechanics in RAGE^-/-^;db/db animals. RAGE inhibition also resulted in improvements to osteocyte density and osteoclast activity. Based on our evidence from longitudinal imaging, we hypothesize that the increased osteoclast activity in FPS-ZM1-treated diabetic animals is responsible for suppressing a marked increase in bone accrual observed in the db/db group between 4-5 months of age. This spike in BV/TV is accompanied by a decline in erosion percentage with no change in mineralization percentage, leading to an increase in steady-state bone percentage. Previous reports have linked decreased bone turnover with impaired material and mechanical properties, likely due to microdamage accumulation^52,53^. RAGE inhibition prevented this increase in steady-state bone percentage in diabetic animals, which likely contributes to the improved matrix material properties and yield load of the db/db+FPS-ZM1 group observed in this study.

As osteoclastogenesis and osteoclast function are suppressed in diabetes^9,10^, our work is the first to show that this osteoclast function can be restored by the inhibition of RAGE after the onset of diabetes. Our data indicate that RAGE inhibition has a significant protective effect on diabetic osteoclast activity, while there trending effects of protection with RAGE deletion. We hypothesize that transient cells such as osteoclasts may be more responsive to abrupt changes in RAGE signaling (e.g. as a therapuetic) than the long-term absence of RAGE. In contrast, osteocytes, which generally have lifespans on the order of decades rather than days, may benefit more from RAGE deletion due to reduced inflammation over the long term. In vitro studies have shown that high levels of glucose can directly impair Receptor Activator of Nuclear Factor kappaB Ligand (RANKL) induced osteoclastogenesis.^54^ At the same time, osteoblasts and osteocytes send signals that increase osteoclastogenesis under high glucose conditions.^15,23^ These contradictory forces suggest than another pathway is shifting the balance to reduce osteoclast activity. We hypothesize that RAGE fills this role. This is supported by a recent single cell RNA sequencing study that revealed a distinct monocyte/macrophage cluster present in T2D mice but not wild-type that exhibited increased expression of AGE-RAGE related signaling pathways.^12^

Our study is also the first to examine the interactive effects of diabetes and RAGE signaling on osteocyte function. Previous in vitro reports have shown that AGEs and high levels of glucose lead to increased osteocyte apoptosis, decreased mechanosensitivity, and altered signaling.^55–57^ Our high-resolution approach to osteocyte imaging complements the existing literature by showing that diabetes not only reduces the number osteocytes per bone volume but also alters the shape of lacunae, making them flatter and less round. Our data shows that RAGE deletion and inhibition increase osteocyte density in diabetic bone, and RAGE deletion improves diabetic lacunar morphology. Previous studies indicate that lacunar morphology influences osteocyte mechanosensitivity,^58^ which, given our findings, suggests that targeting RAGE may not only improve osteocyte numbers but also osteocyte signaling. RAGE deletion and inhibition increased osteocyte density in diabetic but not wild type animals, which suggests that the beneficial effects on osteocyte number stem from the mitigation of inflammatory RAGE signaling in the diabetic state, thereby reducing the frequency of osteocyte apoptosis. Osteocyte senescence could also be playing a role in this phenotype as increased osteocyte senescence has been shown in cortical bone of T2D animals.^20^ These senescent osteocytes show a unique inflammatory profile that may be partially downstream of RAGE signaling.^20^

While other studies have characterized the nanoscale matrix mechanics of the T2D bone phenotype, including one study that showed improved nanoindentation outcomes with RAGE deletion in a high-fat diet mouse model^59^, to our knowledge this is the first study to show that nanoscale matrix mechanics can be altered by pharmacological RAGE inhibition. Our results show that diabetes increases indentation modulus and hardness in mouse trabecular bone, which generally align with previous nanoindentation studies performed in the cortical bone of humans and mice.^52,60,61^ However, indentation viscosity is not reported in these studies and, in our opinion, tends to be underreported in nanoindentation studies of bone. Previous reports show that viscoelastic properties are predictive of bone fracture.^62,63^ Studies have also shown that indentation viscosity correlates with indentation modulus and hardness measurements, highlighting the interplay between the organic and mineral in bone mechanics.^64,65^ In this study, diabetes reduced indentation viscosity, which is consistent with prior reports of increased indentation distance in TallyHo mice^16^ and patients with T2D^66^. Both RAGE deletion and inhibition increased indentation viscosity in diabetic animals but did not affect indentation viscosity in wild-type animals. This is supported by significant ANOVA interaction terms in both models. In this study, RAGE deletion and inhibition improved AGE concentration only in diabetic animals, which is why we hypothesize that this diabetes-dependent action of RAGE on indentation viscosity is driven by improvements to bone composition.

At the whole-bone scale, RAGE deletion protected against reduced stiffness, yield load, ultimate load, work to fracture, and cortical thickness in diabetic animals. However, RAGE inhibition only produced improvements to yield load in diabetic animals. Even though RAGE inhibition in diabetic animals improved bone cell function, trabecular microarchitecture, and matrix mechanics in the vertebra, the treatment used in this study did not replicate the profound improvements to cortical whole-bone mechanics consistent with RAGE deletion. These results align with prior work that shows improvements to diabetic bone with RAGE deletion in both db/db and high-fat diet models^28,59^ and no effect on bone mechanics with Azeliragon (another RAGE inhibitor) treatment in middle-aged mice^67^. However, RAGE inhibition via FPS-ZM1 has shown mediating effects on inflammation in bone marrow mesenchymal stem cells in a high glucose environment.^68^ FPS-ZM1 has also reduced osteocyte inflammatory cytokine release and apoptosis in vitro.^69^ These studies highlight the possibility of reducing chronic NF-κB inflammation in the bone microenvironment with FPS-ZM1 administration. Further research utilizing a longer duration of treatment may be necessary to investigate whether RAGE inhibition can improve whole bone fracture mechanics given our promising data on nano-scale mechanical properties and the existing literature documenting the anti-inflammatory effects of FPS-ZM1.

There are several limitations within this study that should be noted. One, the use of leptin receptor deficient mice as a model for type 2 diabetes. Leptin receptor signaling in osteocytes has a role in the development of cortical bone consolidation^59^ and may participate in on bone cell homeostasis. Additionally, this study was only performed in female mice so differences due to sex were not addressed. Nanoindentation was performed on MMA embedded dehydrated samples, neglecting the important effects of hydration and nanoporosity on nanoscale mechanical properties^38^, though all samples received the same treatment and are therefore comparable within this study. Finally, the measure of AGEs with the total fluorometric assay does not measure the full concentration of AGEs within the bone because non-fluorescent AGEs are not being quantified, though fluorescent AGEs have been shown to correlate with total AGE concentrations in human tissue.^70^

## Acknowledgements

We would like to acknowledge the Washington University in St Louis Musculoskeletal Research Center for their help and expertise, specifically Michael Brodt and Matthew Silva, PhD. Additionally, we would like to thank Rachana Vaidya, PhD for her insights. This work was conducted with funding support from National Institute of Health: R01AR074441, R01AR077678, R21AR081517, T32DK108742, S10OD028573, and P30AR074992. The authors are grateful for the generous gift of the RAGE-null mice from Professor Ann Marie Schmidt of New York University

## Author Contributions

TH- Formal analysis, Investigation, Visualization, Writing; KSB- Conceptualization, Formal analysis, Investigation, Methodology, Validation, Visualization, Writing, Project administration; REW- Investigation, Review, Editing; SR- Investigation, Methodology; KMF- Methodology; SYT- Resources, Supervision, Funding, Writing, Review, Editing.

