## Supplementary material for "Genetic and pharmacologic modulation of RAGE rescues the diabetes-mediated impairments to bone at multiple length scales": S1

Supplementary materials

S1. Viscoelastic Plastic Analysis Curve Fit MATLAB code and example curves

%Oliver Pharr and Viscoelastic Plastic Analysis

%Method based on viscoelastic plastic analysis from Rodriguez-Florez et al

%Variables that change depending on loading rate:

%holding period time: tr<t<tc

prompt="What is the max load for the experiment in uN?"

max=input(prompt);

%Assuming Steady-State Creep:

%hcreep(t)=((k*tr)^(1/2)/(a_3*nq)^(1/2))(t-t1)+h(t1)

tc=30;

tr=12.5;

t1=tr+tc/6;

%k is the loading rate: Pmax/tr: 800uN/s for 10000N tests, 80N/s for 1000N

%tests

k=(max/tr)*10^-6; %N/s

tiledlayout(7,3)

%import and organize data

%% Import data from text file

% Script for importing data from the following text file:

%

%filename: C:\Users\brozk\OneDrive - Washington University in St. Louis\Nanoindentation Analysis\Raw data\1962\1000\1962_1000_Position 0_00000 LC_20240411_1853.txt

%

% Auto-generated by MATLAB on 18-Jun-2024 15:01:08

files=dir('*.txt');

for i=1:length(files)

%% Set up the Import Options and import the data

opts = delimitedTextImportOptions("NumVariables", 5);

% Specify range and delimiter

opts.DataLines = [5, Inf];

opts.Delimiter = "\t";

% Specify column names and types

opts.VariableNames = ["Depthnm", "LoadN", "Times", "DepthV", "LoadV"];

opts.VariableTypes = ["double", "double", "double", "double", "double"];

% Specify file level properties

opts.ExtraColumnsRule = "ignore";

opts.EmptyLineRule = "read";

% Import the data

p11= readtable(files(i).name, opts);

%% Clear temporary variables

clear opts

%tc: creep time, tc=30s

%first delete the approach and nans

a=find(p11.Times<5);

p11(a,:)=[]; %Remove the approach time

p11=rmmissing(p11); %Remove nans

%Setting time at beginning equal to t0, approach= 5s

p11.Times=p11.Times-5;

%Assuming Linear creep: hcreep(t)=((k*tr)^(1/2)/(a_3*nq)^(1/2))(t-tr)+hload(tr)

%alphas=a are the geometric constants for a berkovich tip

a_3=4.4;

%Non-linear least quatres curve-fit function

%find C aka h(t1)

a=find(p11.Times>=t1);

b=find(p11.Times>=tr+tc);

C=p11.Depthnm(a(1))*1e-9;

%FINDING Nq

%ydata=(((k.*tr).^(1/2))./((a_3.*x(1)).^(1/2)))*(xdata-t1)+C

ydata=p11.Depthnm(a(1):b(1)).*1e-9;

xdata=p11.Times(a(1):b(1));

fun=@(x,xdata)x*(xdata-t1)+C	% ASSUMING THAT h(tr===h(t1))

x0=[100];

options = optimoptions('lsqcurvefit','Algorithm','levenberg-marquardt');

[x,resnorm,residual,exitflag,output]=lsqcurvefit(fun,x0,xdata,ydata);

nexttile

plot(xdata,ydata,'ko',xdata,fun(x,xdata),'b-')

s=x;

nq=(1/a_3)*((k*tr)^(0.5)/s)^2;

vars={'xdata','ydata','a','b','C','t1','x','resnorm','residual','exitflag','output'};

clear vars;

NQall(i,1)=nq;

end

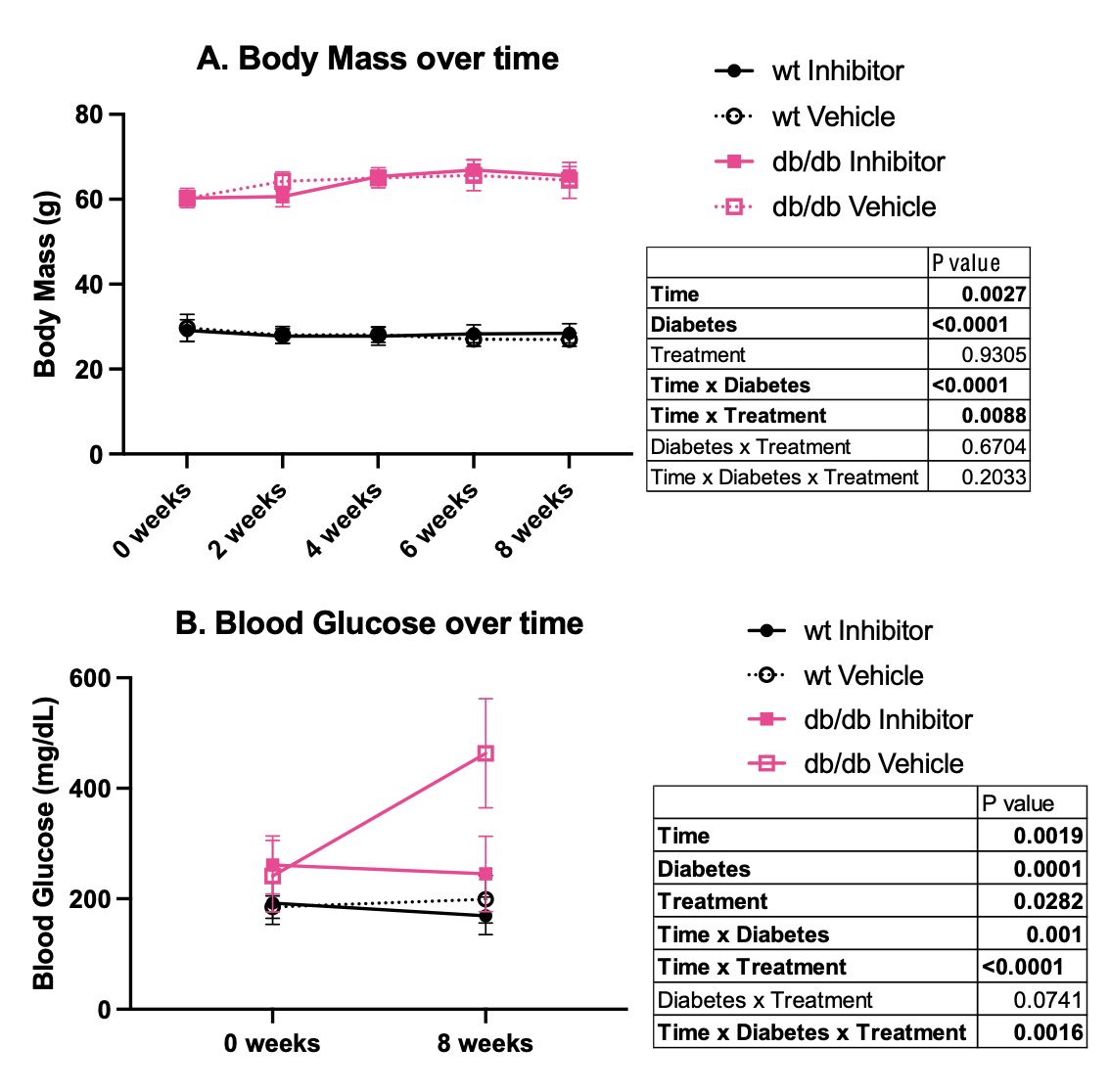

S2. RAGE inhibition with FPS-ZM1 prevented blood glucose increase in db/db animals at 5 months of age (8 weeks of treatment) compared to 3 months (0 weeks of treatment). (A) Body mass was consistent across 3 months to 5 months of age in both wt and db/db animals, with no obvious effect of FPS-ZM1 treatment over time. (B) Blood glucose increased over time in the db/db vehicle group, but not in the db/db inhibitor group.
